# Interpreting sensory and cognitive signals in the cortical reading network

**DOI:** 10.1101/2021.09.14.460238

**Authors:** Garikoitz Lerma-Usabiaga, Rosemary Le, Chen Gafni, Michal Ben-Shachar, Brian Wandell

## Abstract

Voxels in the visual cortex contain neurons that are responsive to small regions of the visual field; these regions can be quantified using population receptive field (pRF) analysis. We measured pRFs using several types of patterns: words, visually matched false-fonts, and visually dissimilar checker patterns. In visual field maps pRF estimates using words and false-fonts are similar, but pRF estimates using checkers differ substantially. The pattern of results in the contiguous ventral occipito-temporal reading circuitry is different: word pRF estimates differ from false-fonts and checkers. These findings were replicated at two research sites. We explain these results with a qualitative model, where the response differences in the visual field maps arise from sensory signals, and the differences in the reading circuitry arise from the integration of sensory and cognitive signals.

## Introduction

Many important tasks, such as reading, require the coordination of sensory and cognitive processing. Regions of ventral occipito-temporal (VOT) cortex, close to the visual field maps, are part of the neural circuitry that is engaged by reading (Dehaene and Cohen 2011; Kravitz et al. 2008; Price and Devlin 2011; Rauschecker et al. 2012; Wandell et al. 2012). Multiple investigators have shown reproducible reading-related activations using functional magnetic resonance imaging (fMRI), electrocorticography (ECoG), and electroencephalogram (EEG) experiments across individuals and orthographies (Baker et al. 2007; Bolger et al. 2005; Glezer and Riesenhuber 2013; Krafnick et al. 2016; Woolnough et al. 2021; Lochy et al. 2018; Nobre et al. 1994). We refer to the regions of the VOT that are more responsive to words than to other stimulus categories as the VOT reading circuitry (VOTRC).

In previous work (Le et al. 2017), we used fMRI and the population receptive field (pRF; Dumoulin and Wandell 2008) analysis to characterize the visual responses in the VOTRC. A pRF describes the region of visual space in which a stimulus elicits a response from a voxel. Combining the pRFs of nearby voxels into a region of interest (ROI), we obtain the field of view (FOV) of the ROI. The FOV defines the portion of the visual field that reliably evokes a response in any of the ROI voxels. Le et al. (2017) showed that the FOV of the VOTRC in the left hemisphere is largely confined to the central portion of the visual field, with a right visual field bias. Furthermore, the FOV was reported to be stimulus-dependent, differing when contrast patterns comprised words versus checkers.

This finding can be explained by at least two types of neural models. The first model is that responses in VOTRC voxels are dominated by sensory input, and this input depends on the stimulus. Changing the stimulus from checkers to words shifts the sensory input so that the VOTRC receives signals from different regions of visual space. A second model is that the signals reaching the VOTRC include a large contribution from non-visual, cognitive regions. In this case, using the pRF model to analyze the VOTRC responses incorrectly assigns differences in the BOLD response to changes in receptive field center locations and sizes.

To understand the origin of this stimulus dependency in the VOTRC, we expand on the prior reports in several ways. First, we quantify the stimulus-dependency in the ventral visual field maps (V1, V2, V3, hV4, VO-1) whose signals are delivered to the VOTRC. A small stimulus-dependent effect is measured as early as in V1 and becomes more pronounced in every subsequent map. The stimulus-dependent differences in pRF center location are systematic: compared to checker stimuli, the word stimuli generally produce estimates that are as much as 3-4 deg closer to the fovea and fall along radial visual field lines. We further observe stimulus-dependent differences in pRF center positions in experiments with false fonts and also in experiments with bilingual subjects who viewed several stimuli, including Hebrew script. In all experiments participants were instructed to perform a simple fixation task and were not asked to read.

The results reveal substantial stimulus-dependent differences in pRF centers in the visual field maps. Stimulus-dependent differences are also present in VOTRC, but they have a slightly different character. Comparisons between words and false fonts show significant differences between the signals in the VOTRC and a relatively posterior visual field map, VO-1. Specifically, VO-1 responds similarly to words and false fonts, whereas the VOTRC responses differ between false fonts and words. In VOTRC, we suggest that the signals are a combination of sensory and cognitive terms. We introduce a model to account for both the findings in the field maps and the VOTRC.

## Results

### pRF centers are closer to the fovea when measured with words than checkers

We begin by comparing pRF eccentricity values in the VOTRC and the ventral visual maps measured using word and checker stimuli. Our stimulus consisted of bars that swept the 15 degree radius visual field, going back and forth along the vertical direction, the horizontal direction, 45 degrees and −45 degrees (see Le et al. 2017). The word and checker stimuli were exposed within the traveling bars. The estimated visual field eccentricity depends on the stimulus substantially in the VOT and as well as in portions of the parietal cortex (Figure 1A), such that the centers of pRFs measured with word stimuli are more foveal than those measured with checker stimuli. Specifically, VOT regions have an estimated eccentricity between 4-7 deg (yellow-green) when measured with checkers, and an estimated eccentricity of 2-5 deg (orange-red) when measured with words. The same effect is seen in the intraparietal sulcus (IPS; see Figure S1A-B).

**Figure 1.**
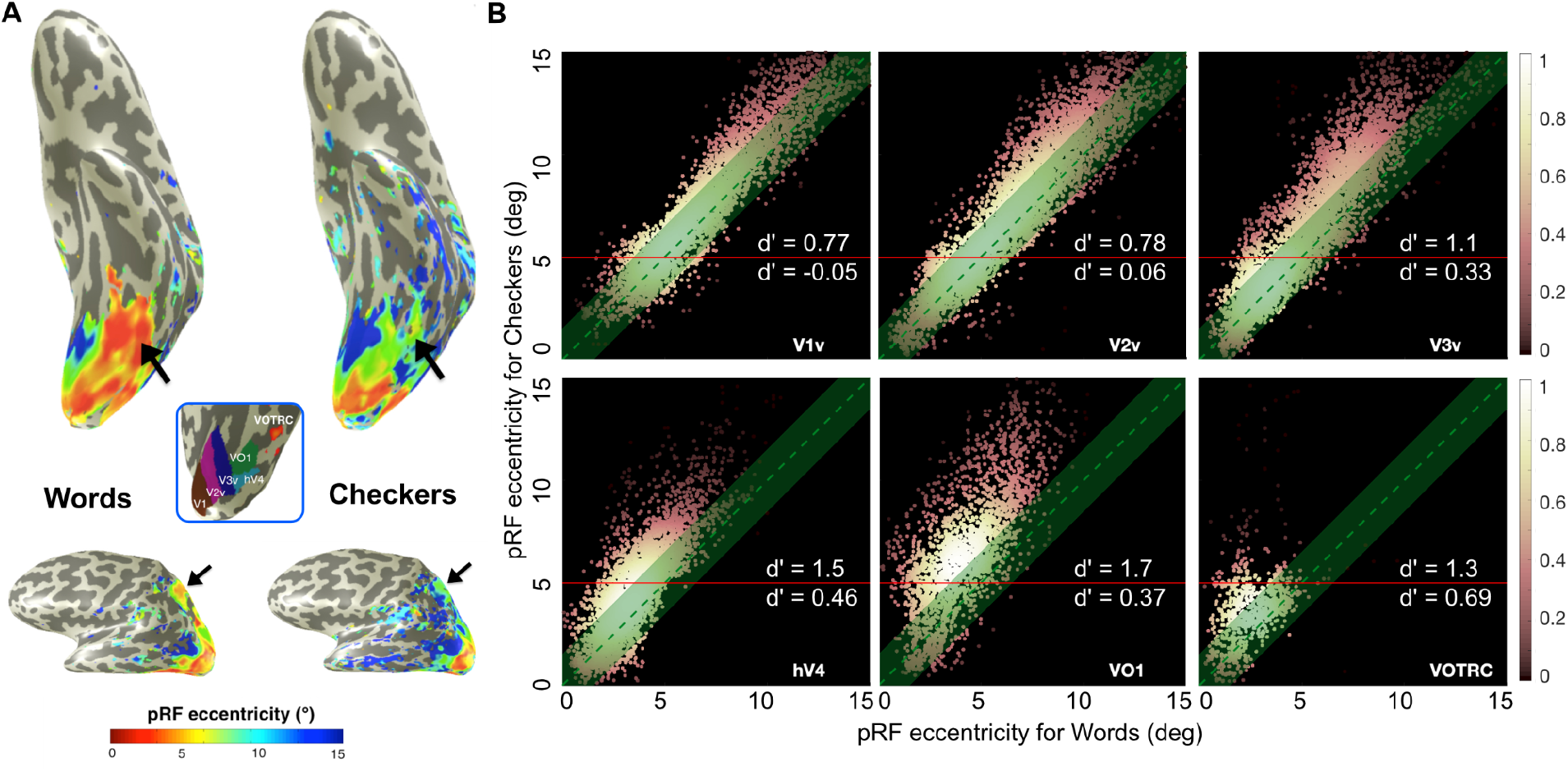
The centers of pRFs measured with word stimuli are more foveal than those measured with checker stimuli. **A.** Top: Two measurements from the ventral surface of the left hemisphere of a typical subject (#20) are shown. Bottom: A lateral view of the same measurements. The four arrows point to regions with considerable stimulus-dependency of the pRF centers. The color map represents the pRF eccentricity (deg) estimated in response to words (left) and checkers (right). Blue inset: the typical positions of the ventral visual field maps (V1v, V2v, V3v, hV4, VO1) and the VOTRC. **B.** Color coded scatter plots comparing the pRF center eccentricities between words and checkers. The color shows the relative density of points within a square region (1 × 1 deg) centered on the point. Each panel shows data for one of the left ventral visual field maps or the VOTRC. For many voxels, the pRF center eccentricity is closer to the fovea when measured with words than checkers (points above the diagonal). The green band shows the region within 1.5 deg of the identity line (representing stimulus invariance). The stimulus-dependent eccentricity difference is largest for voxels with eccentricities near the periphery (5-15 deg), above the horizontal red line. The effect size above and below the 5 deg line is measured by the d’ values inset in each panel. The data are based on voxels in which the pRF model explains at least 20% of the variance. The variance explained using words and checkers was similar across the visual maps and VOTRC, with a slight advantage to checkers in the early maps and slight advantage to words in the VOTRC (Figure S1C).

There are also stimulus-dependent differences in pRF eccentricity estimates in the visual field maps (V1, V2, V3, hV4, VO1) and the VOTRC. These differences can be visualized in scatter plots that compare estimated eccentricity of the pRF centers in response to words and checker stimuli in individual voxels (Figure 1B). The plots show that there is a small stimulus-dependent effect in V1 and V2, and an increasingly large effect in V3, hV4, VO1 and VOTRC. The pattern of the scatter plots is quite similar in V1-VO1, so we discuss those first. We will discuss the findings in VOTRC separately below.

The pRF center in V1-VO1 tends to be closer to the fovea when measuring with words compared to checkers. The stimulus-dependent eccentricity shifts are substantial when measured in voxels whose pRF centers are from 5 to 12 deg eccentricity (referred as near periphery from now on). We quantify the effect in two ways. First, the size of stimulus-referred shift is on the order of 1-3 deg. Second, we compare the size of the shift to the standard deviation of the measurements (d’, or Cohen’s d). We separate the voxels in two groups, parafoveal voxels (when checkers’s eccentricity is less than 5 deg) and near periphery, indicated by a horizontal red line in Figure 1B. For parafoveal V1 and V2, the d’ is about zero. The stimulus-dependent shift increases in V3, hV4 and VO1, reaching about d’=0.4. For near periphery eccentricities, the stimulus-dependent eccentricity effect size in V1 is d’=0.7, increasing to d’=1.7 in VO1 (Figure 1B).

The VOTRC scatter plot seems qualitatively different from the others. The main reason is that even when measured with checkers, there are very few voxels with a center in the near periphery. Thus, the stimulus-dependent effect is measured using mainly the voxels in the parafoveal region which limits the potential effect size. For this reason, the linear trend with eccentricity that is easily observed in V1-VO1 does not seem appropriate for the VOTRC data. The main finding remains: the pRF eccentricity estimates are closer to the fovea when measured with words compared to checkers (near periphery d’=1.3, parafoveal d’=0.7).

We are measuring the effect using the parameters of the pRF model, and thus we should understand how accurately the model predicts the measured responses. There were no remarkable differences in variance explained between the two conditions, words and checkers (see Figure S1a). The level of variance explained is comparable to the variance explained when replicating the data. That is, participants underwent two measurements and the first scan was used as a model for the second: the median variance-explained ranges from approximately 60% in V1, declining through the field maps to 10% in VOTRC. There are various ways to parameterize these effects, which is why we present the full scatter plots and additional quantitative analyses in Figure 1.

In the scatter plots we show voxels for which the pRF model explains more than 20% of the response variance for both conditions. This criterion with respect to variance explained varies across studies (Gomez et al. 2018; Benson et al. 2018; Infanti and Schwarzkopf 2020; Harvey and Dumoulin 2011; Alvarez et al. 2015). Our measurements are shared, and thus anyone who cares to explore the data at different threshold levels is welcome to test both the data and our software methods. Overall, the findings generalize across many different thresholding strategies (see Figure S1B).

Next we examine the change in the pRF center angle within the visual field. To simplify the visualization, we translated and rotated the checker center position to the horizontal axis (15 deg position), and rotated and translated the two pRF center estimates by the same amount (Figure 2A). The distribution of differences in eccentricity (represented by Δ in Figure 2A), is plotted for the near periphery (Figure 2B top) and parafovea (Figure 2B bottom). In the near periphery the stimulus-dependent eccentricity effect increases from posterior to anterior regions. In the parafovea, the effect is smaller but present.

**Figure 2.**
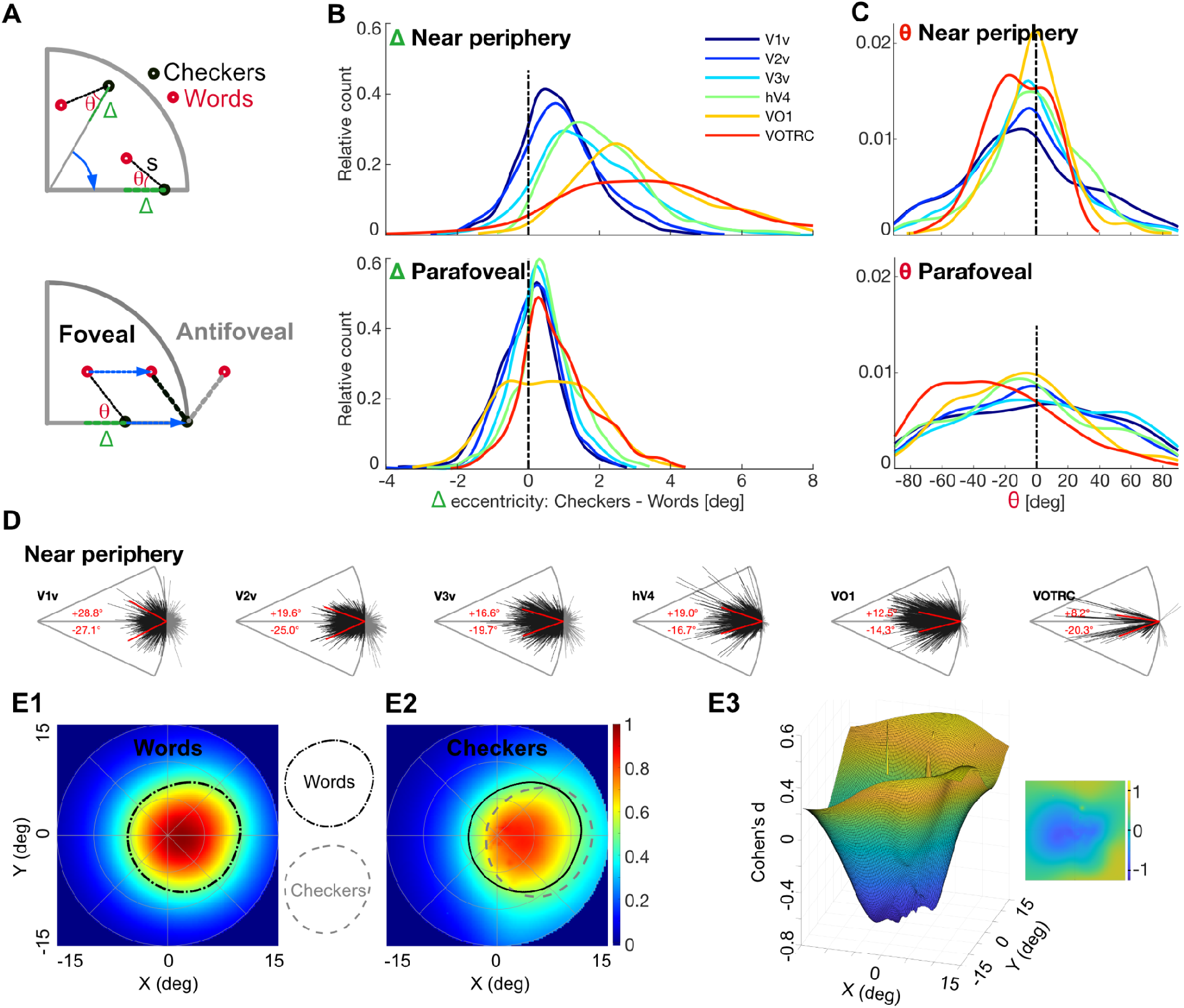
The word pRFs centers are more foveal along radial visual field lines than checker’s center. **A.** The parameterization of the differences between word and checker pRF centers is illustrated. The two measurements are shown as red (word) and black (checker) dots on a visual field representation (upper right quarter is shown). The two measurements are rotated and translated (blue arrows) so that the pRF center of the checker data is on the same location in the x-axis. This enables us to compare voxels with many different centers. The eccentricity difference is represented by the green segment (Δ). The radiality is measured by the angle between the x-axis and the segment (s) connecting the two pRF centers (Θ). **B.** The distribution of Δ (stimulus-dependent eccentricity difference) for voxels in the near periphery (top) and parafovea (bottom). The curves are distributions for the ventral field maps and the VOTRC. All areas under the distributions are equal to 1. **C.** The distribution of Θ, organized as in **B**. **D.** The x-axis aligned segments (s) between the two pRF centers, expressing the angle (Θ) between the checker pRF center and the fovea. Beyond V1 the segments in the near periphery cluster around the x-axis, revealing a tendency for the pRF center to shift towards the fovea. The red lines are the median of the positive and negative angles. Only near-periphery is shown as the parafoveal results show less radiality (see 2C). **E.** Combining the pRFs we can specify the field of view (FOV) available to the voxels in a region of interest. These FOV plots show the mean pRF amplitude across subjects at each location in the visual field from the voxels in the VOTRC (the half contour of the FOV is represented with the dash-dot line in E1 and the gray dashed line in E2). Measured with words (E1) the FOV is concentrated near the fovea; measured with checkers (E2) the FOV is displaced to the right. The black line in E2 is a collection of 25 bootstrapped with replacement half contour lines. E3 shows the effect size (d’) of the pixel-wise difference between the word and checkers FOVs. The inset shows the same data in a 2D view for clarity.

To check a stimulus-dependent angle effect, we measured the density distribution of the angle difference (represented by Θ in Figure 2A). If the pRF center displacement tends to be radial (towards the fovea), the distributions Θ should be concentrated around zero. In the near periphery we see this concentration (Figure 2C top), particularly in the VOTRC. In the parafovea the effect is not clearly present (Figure 2C bottom). This suggests that pRF center changes for parafoveal voxels are not consistently more foveal. Figure 2D shows the angles and lengths of the difference segments (s) for all near periphery (top) voxels. The s segments are black and inside the circle when the word pRF center is more foveal than the checker pRF center, and gray and outside the circle otherwise. The concentration near the horizontal axis and towards the fovea increases through the visual field maps and is clear in VOTRC voxels.

### PRF centers measured with false-fonts and words differ in the VOTRC but not in VO1

Next we report pRF measurements made with false-font stimuli. False-font stimuli are visually more similar to words than to checker stimuli, but are not readable. In V1v-V2v-V3v-hV4 we observe no differences or small differences between false-font and word eccentricity values (see Figure S2A-B), but false-fonts vs. checkers follow a pattern of differences similar to words vs. checkers (see Figure S3A-B).

The differences between false-fonts, words and checkers are best illustrated by comparing the results in V01 and VOTRC (see Figure 3). In VO1 (top row), false-fonts are similar to words but different from checkers; in VOTRC (bottom row), false-fonts are different from words but similar to checkers.

**Figure 3.**
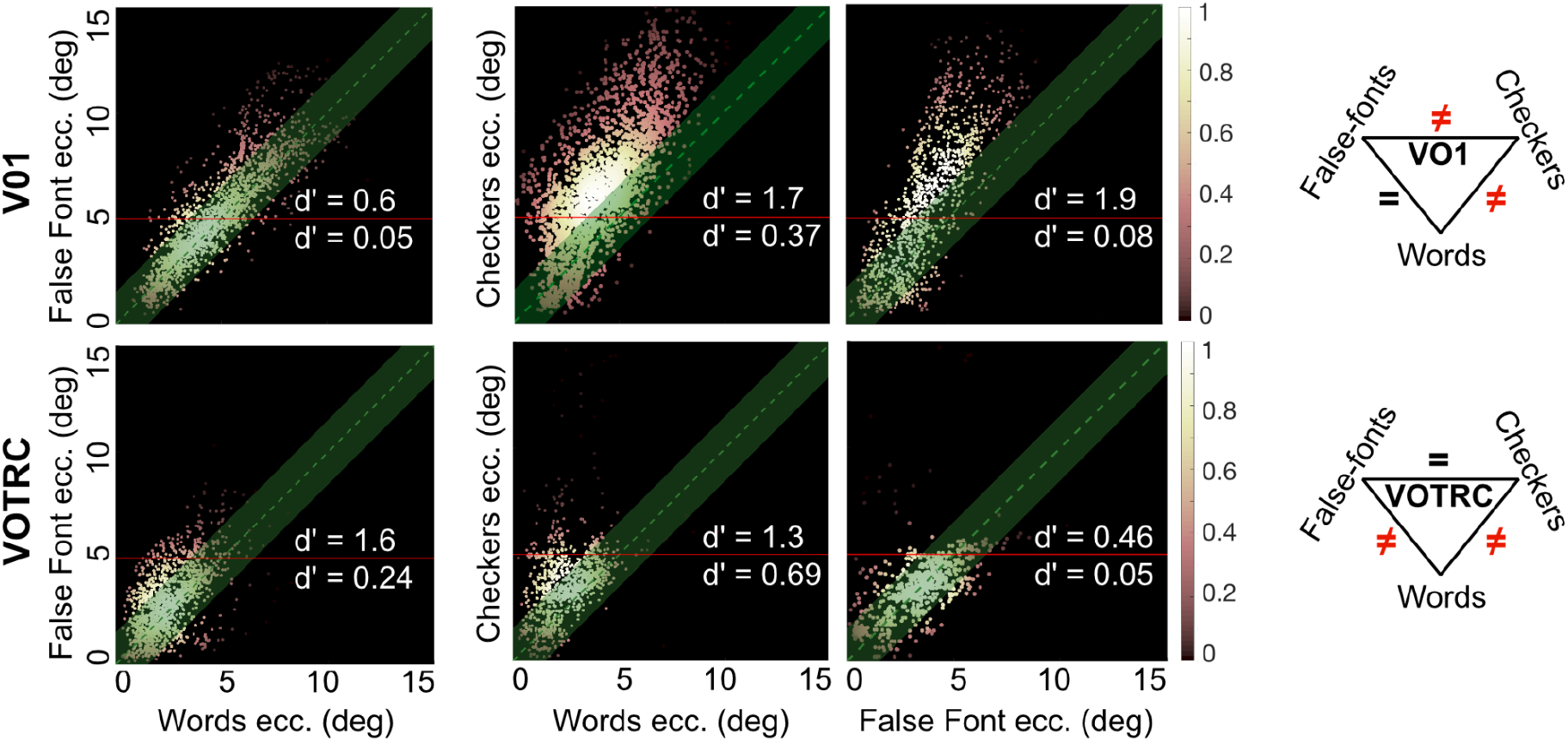
Comparison of the pRF centers for English words, False Fonts, and Checkers. Words versus false-fonts eccentricity scatterplots in VO1 (top) and VOTRC (bottom). Each scatterplot compares two of the three stimuli: Words vs False Fonts (left), Words vs. Checkers (middle) and False-fonts vs. Checkers (right). The stimulus-dependence changes between VO-1 and VOTRC. In the diagram: in VO-1 the Words and False Fonts, similar stimuli, are equivalent and both differ from Checkers; in VOTRC the non-Word stimuli both differ from Words. All voxels with variance explained more than 20%. Eccentricity values for variance explained at 5% in Figures S1B–S2B–S3B, and comparison of variance explained in Figures S1C–S2C–S3C. Scatterplot details as in Figure 1B.

### VOTRC FOV differences generalize when measured using different alphabets, MR scanners and displays

Bilingual readers (L1 Hebrew and L2 English; N=13) viewed the same bar stimuli described above, but in this case containing Checkers, Hebrew and English words (these measurements were performed in a different scanner and with a different field of view, see Methods for more details). Hebrew and English written words differ in many aspects, e.g., orthographic (the letters are different), lexical (word identity is different), and in reading direction (Hebrew is read from right to left). The first set of results is confirmatory: the English vs. checkers contrast in the L2-English group replicated the results for this contrast in the L1-English group. We observe the same effect when comparing Hebrew words to Checkers. In both cases, words evoke more foveal responses compared to checkers, and the effect grows increasingly larger from posterior to anterior regions (V1-V3, hV4,VO-1, VOTRC; see Figure S4).

The second set of results shows that eccentricities of word responses in Hebrew and English are the same in visual field maps, except some minor differences in the VOTRC (see Figure S5 and S7). This difference is better observed when showing the pRFs aggregated as the FOV (see Figure 4A). In summary, the FOV for Hebrew is more leftward than the FOV for English, which is itself more leftward than Checkers (Figure S6 shows significant differences in pRFs X coordinate, but not in the Y coordinate).

**Figure 4.**
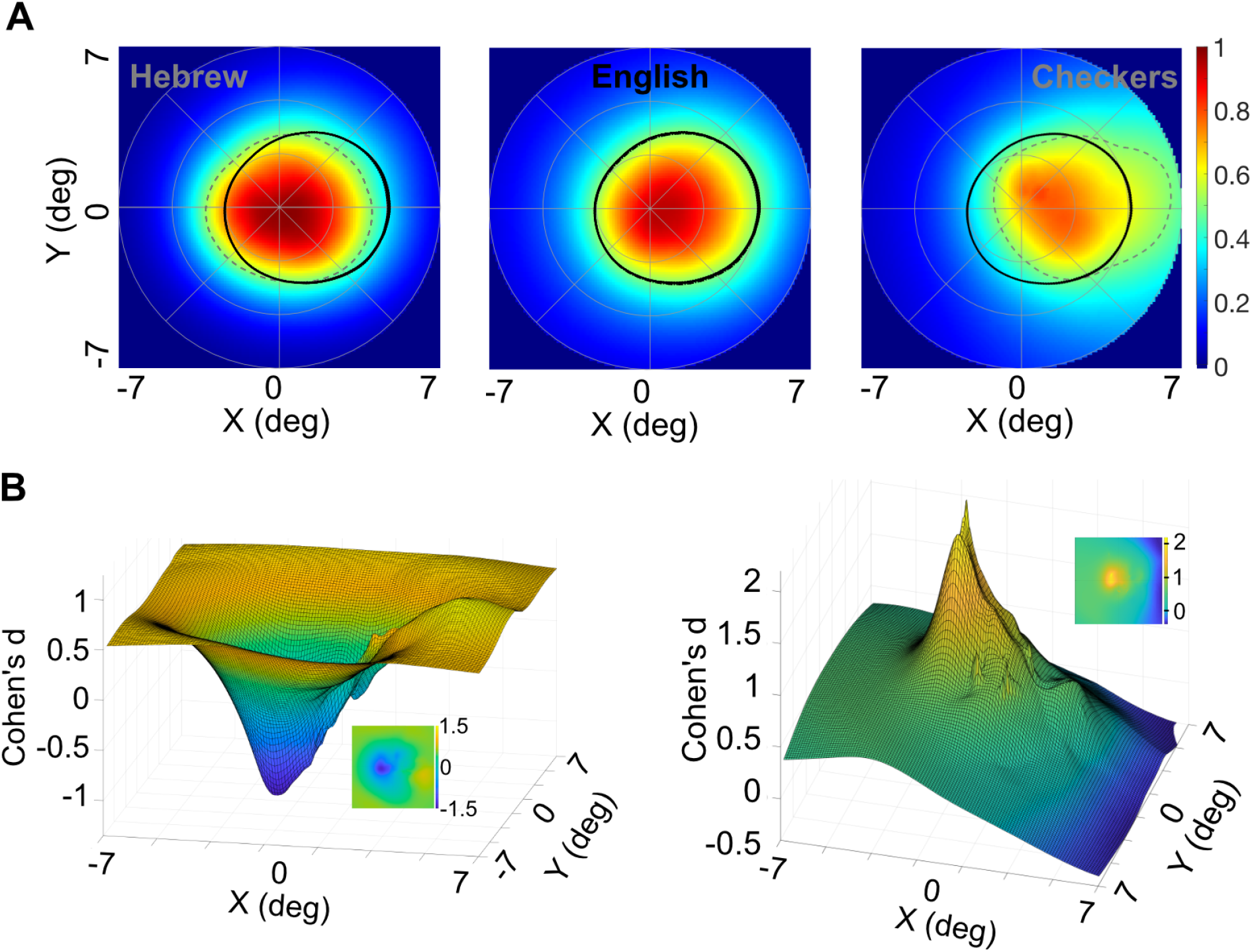
Position sensitivity with English words, Hebrew words and checkers (N = 13). **A**. FOV coverage in VOTRC for Hebrew words (left), English (center) and Checkers (right). Grey dashed lines correspond to the 50% contour line in Hebrew and Checkers. Black lines in all three panels are the same, and correspond to the 50% contour line for English words (50 bootstrapped with replacement lines). **B**. Effect size (Cohen’s d) representation of the difference in FOV for English - Hebrew words (left) and English Words - Checkers (right). The values shown in the mesh (and in flat format in the inset) are the mean of 100 bootstraps with replacement (50 repetitions). The comparisons were paired, so only subjects with enough surviving voxels were used in the comparisons (N=9 for English - Hebrew; N=8 English - Checkers). Minimum and maximum d’ values for English and Hebrew were −1.4 and 0.8, respectively. Minimum and maximum d’ values for English and Checkers were −.4 and 2.2 respectively. In this figure only voxels with more than 20% of the variance explained were selected, see Figures S4B–S5B for scatter plots with 5% variance explained, where more voxels survive and the same results can be observed. This effect was observed in most subjects at the individual level (Figure S7). The difference between English and Hebrew words were clear in the VOTRC, but not in the other visual field maps (Figure S7).

## Discussion

The Introduction contrasts two models to explain the pattern of results reported in VOTRC (Le et al. 2017). In one model, the stimulus-dependent VOTRC responses start in the visual field maps. In this case, each VOTRC voxel receives signals from different parts of visual space. In a second model the VOTRC responses comprise a mixture of sensory and cognitive signals. In this case the pRF model, which attributes the stimulus-dependent changes entirely to sensory input, is incomplete and requires elaboration.

The analysis here shows that there is a large stimulus-dependent effect on the pRF in the visual field maps. Starting as early as V1 or V2 and continuing through VO-1, we find a consistent difference between the responses to checkers compared to both words and false-fonts (Figures 1–3). In the transition into the VOTRC, however, there is yet another stimulus-dependent change in the response. In VO-1, responses to false fonts and words are similar, and both differ from the response to checkers. In VOTRC, responses to false fonts and checkers are similar, and both differ from the responses to words (Figure 3; see triangle diagrams). Moreover, the VO-1 and VOTRC responses to words align, but the VO-1 and VOTRC responses to false fonts do not align (Figure 3; see differences between rows).

Comparing English and Hebrew words, the FOV is very similar in all the visual field maps. There is a small difference in the FOV measured in the VOTRC. The difference is a horizontal shift; the FOV coverage for the Hebrew words is shifted slightly towards the reading direction (left for Hebrew and right for English).

These changes in the response pattern from VO-1 to VOTRC do not appear to be a sensory effect; words and false-fonts have similar visual characteristics as do Hebrew and English. The difference appears to be because of the cognitive differences (word or not, for example reading direction). We suggest that VOTRC responses include a cognitive contribution that distinguishes words as lexical units from false fonts as visual objects (Yeatman and White 2021; Lerma-Usabiaga et al. 2018), and the differences between English and Hebrew words arise from cognitive differences related to the different languages.

The region in visual space where a stimulus evokes a neuron’s response is the neuron’s receptive field. The point spread function is a complementary measure: the region of cortex that responds to a point stimulus. Point spread functions in the visual cortex of human and macaque, measured both with fMRI and electrophysiology, have a half maximum spatial extent of about 5 millimeters (Engel et al. 1997; Harvey and Dumoulin 2011; Palmer et al. 2012). This suggests several ways in which the neural pathways can alter the receptive field of a neuron or an fMRI voxel.

One mechanism for altering the receptive field is through adjusting the signals between different cortical regions. Feedback connections between different regions may be a means of regulating the connection strengths. Electrophysiological evidence demonstrates that the receptive field properties in any given map result from interactions of neurons both within that map and other maps. Nurminen et al. (2018) show that blocking the feedback from V2 to V1 alters the size of the V1 receptive field. Kirchberger et al. (2021) showed how the feedback signals to V1 impact the ability to discriminate an object from its background. A second mechanism for altering the receptive field is at the synaptic level. El-Boustani et al. (2018) showed how the receptive field of a V1 neuron can be altered by spike timing-dependent plasticity changes at the synaptic inputs. Stimulus dependent variations in neural responses have been reported in other contexts as well (Gutnisky and Dragoi 2008; Bányai et al. 2019).

Some of these changes can be attributed to properties of the stimulus or stimulus-induced changes. In addition, experimental data support the idea that tasks modify the receptive field in both human and animal models. For example, covert attention influences the receptive field of a neuron or an fMRI voxel (van Es et al. 2018; Kay et al. 2015; Zhang and Kay 2020; Treue and Martinez-Trujillo 2012).

Figure 5A sketches how the anatomical connections in the visual cortex might enable a stimulus-dependent shift in the pRF through re-weighting input connections. In this diagram, two inputs (red, blue) with different spatial contrast are presented at the same retinal location. One stimulus (blue) preferentially excites neurons that are closer to the fovea. Consequently, the spatial distribution of activity in V1 between the stimuli will differ slightly. The transmission from V1 to V2 is shown as amplifying this separation; the feedback loops shown as gray arrows may be a mechanism that governs this weighting. The size of the stimulus-dependent change in connection strength is shown as being small in V1 and V2, and the separation is amplified as the signal propagates through the maps. It is also likely that the effects are largest for neurons that represent the peripheral visual field which have the largest receptive field size. This computational architecture enables a stimulus-dependent change in the pRF that is consistent with our observations.

**Figure 5.**
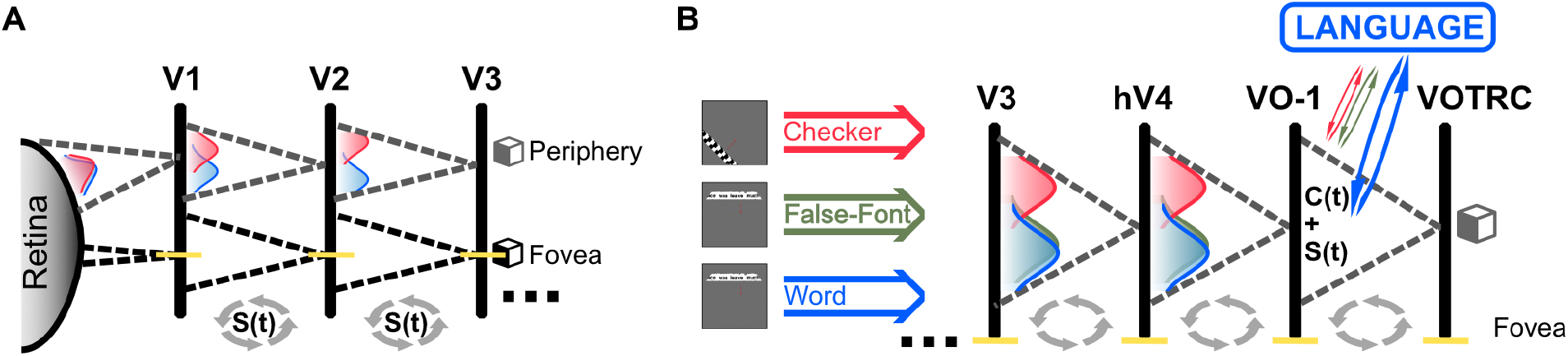
Model for interpreting sensory and cognitive signals in the cortical reading network. **A.** The response of a voxel depends on the structural and functional neuronal architecture. The dashed lines represent the structural connectivity of a voxel in V1-3, representing all the possible input neurons that could drive a response in the connected voxel. The comparison between the black and gray dashed lines between V1 and the retina illustrate how the RF of a V1 voxel will be narrower in the fovea and larger in the periphery. Within the anatomical constraints, the system has the ability to adapt. The signal is present at the input of all connected neurons in the voxel, and depending on the spatial characteristics of the stimuli, the signal will choose one neuronal path or another. We represent the likelihood of those functionally chosen paths with the colored Gaussian probability distributions. In this 1D example, the position of the Gaussian on top of the black continuous line represents the eccentricity of the pRF, i.e.: the gray voxel in V3, depending on the spatial characteristics of the stimulus, will be processing a pRF with different eccentricities. Gray arrows in **A** and **B** indicate that the RF properties in any given map result from sensory signal S(t) interactions of neurons both within that map and other maps. **B.** The signal for each type of stimulus is represented with different colors (red for checkers, green for false-fonts and blue for words). In our example, a hV4 voxel implements the red Gaussian (checkers) separated from the green and blue Gaussians (false-fonts and words respectively, which are very similar visually). The VOTRC voxel is different: it integrates the incoming sensory signals from VO-1 and the cognitive signals from the language network. Due to this integration, the cognitive signals from the language regions are capable of separating the false-fonts from the words. Blue, red and green arrows from the Language network represent the cognitive signal C(t) capable of differentiating the words from other stimuli. VOTRC combines sensory S(t) and cognitive C(t) signals, and limits the interpretability of the BOLD signal with the classical pRF model.

The sensory model can explain responses in the visual maps, but not in the VOTRC. The responses to words and false-fonts are similar through all of the visual field maps, but the response properties differ between VO-1 and VOTRC. This difference may arise from the cortical systems that identify the stimulus as a word and deliver cognitive signals that combine with the signals from the visual field maps. Figure 5B shows cognitive signals arriving to VOTRC that differ substantially when the stimuli are words compared to checkers and false fonts. At this point, we do not have a quantitative model of the cognitive signal. We can only report that it differs between words and false-fonts.

Cognitive signals in the VOTRC are not part of the pRF model which makes a pure input-referred interpretation of the responses. If there is a substantial cognitive signal, its properties are such that the pRF interpretation of the word responses continue to be relatively foveal, while its impact on the false font signals makes them more similar to the checker stimuli.

Could this cognitive signal also explain the differences in the visual field maps? Because the responses in the visual field maps do not distinguish between false fonts and words, we do not think it is part of that system. That is why we favor the idea that stimulus-dependence arises from processing within the maps themselves (Figure 5A). We speculate that the cognitive signals that separate responses by type become significant as the signals enter the VOTRC (Figure 5B).

Given the extensive network of white matter connections in the brain, it seems nearly certain that multiple regions will contribute to the responses in association areas. Experiments supporting this idea in ventral occipito-temporal cortex were previously described by (Kay and Yeatman 2017), who report task-effects in VOTRC. The VOTRC is quite likely to be part of a larger language and attentional network (Chen et al. 2019), and other nearby ventral cortex regions are likely to be part of brain-wide networks as well.

It is worth noting that our participants were not asked to read. They were instructed to perform a simple fixation task. Yet, we find a difference between false-fonts and words in VOTRC. The participants always notice that the stimuli sometimes includes words. The VOTRC response differs between words and false-fonts, and this distinction may be based on cognitive signals from other brain regions. This possibility is supported by measurements of the temporal dynamics of the VOT response. First, a number of investigators observed that responses in the inferior frontal gyrus take place earlier in time than responses in the VOTRC (Tarkiainen et al. 2002; Pammer et al. 2004; Cornelissen et al. 2009; Woolnough et al. 2021). Also, there is a 100 ms delay between different regions within the VOTRC (Boring et al. 2021), which is far longer than the neural transmission time. These findings support the idea that VOTRC processing is at the nexus of sensory and cognitive signals (Price and Devlin 2011; Carreiras et al. 2014; Yeatman and White 2021; Lerma-Usabiaga et al. 2018).

We conclude that there is a substantial stimulus-dependent pRF eccentricity shift in regions that represent the near periphery in the early visual field maps. The pRF measured with stimuli comprising thin lines (text and false font) is more foveal than the pRF measured using checker patterns. The shift is clear in voxels representing the periphery, and there is either a small or no shift for voxels in the central few degrees. The stimulus-dependent eccentricity shift was observed using different alphabets (Hebrew and English) and acquisition parameters at scanning centers. Experiments with additional stimulus manipulations may clarify the critical stimulus parameters that cause these shifts; quantitative circuit modeling may be helpful to understand the difference between foveal and near-periphery responses.

Responses in the VOTRC require a different explanation. First, nearly all of the pRF centers are within the central five degrees. Second, the responses to false fonts and checkers are more similar, not the responses to false fonts and letters. To understand these responses, we suspect it is necessary to extend the purely sensory pRF model to account for the presence of significant cognitive signals that define the stimulus category. This goal can be achieved by developing a model that includes an explicit cognitive signal, and how this signal alters the BOLD response.

## Acknowledgments

This project received funding from the European Union’s Horizon 2020 research and innovation programme under the Marie Sklodowska-Curie grant agreement No 795807 to G.L.-U and from the Binational Science Foundation, grant number 2011314 awarded to B.W and M.B-S.

## Author contributions

G.L-U, R.L., M.B-S., and B.W. Conceptualization, Methodology, Writing G.L-U, R.L., and B.W. Software, Validation, Formal Analysis, Resources, C.G., R.L. Investigation, B.W., M. B-S. G. L-U. Funding and Supervision.

## Methods

### Subjects

Twenty right-handed subjects (12 females; median age 24 years, range 20–35 years) were scanned at the Center for Neurobiological Imaging (CNI) at Stanford University. All subjects gave informed consent, were right-handed native English speakers, and had normal or corrected-to-normal vision. The scanning was approved by the Institutional Review Board at Stanford University.

Fourteen right-handed subjects (10 females; median age 25 years, range 18-37) were scanned at the Strauss Center for Computational Neuroimaging at Tel Aviv University. All subjects gave informed consent, were right-handed, and had normal or corrected-to-normal vision. These subjects were native Hebrew speakers and could read both Hebrew and English. The scanning was approved by the ethics committees of the Sheba Medical Center and Tel Aviv University.

### Neuroimaging

Data acquired at the CNI used a 3T General Electric MR 750 scanner with either a Nova 16- or 32-channel head coil. The Nova 16-channel is identical to the Nova 32-channel coil, but with the front half of the coil removed to allow for an unobstructed field of view. Data acquired at the Strauss Center for Computational Neuroimaging at Tel Aviv University used a 3T Siemens Magnetom Prisma scanner with a Siemens 20-channel head coil.

#### Anatomical

Anatomical data acquired at the CNI used the 32-channel coil with a 3D Fast SPGR scan (166 sagittal slices, resolution 0.9375 x 0.9375 x 1mm). For each subject, 1-3 anatomical volumes were acquired, averaged, and resampled to 1mm isotropic voxels. Anatomical data acquired at the Strauss Center used the 20-channel coil with a 3D Fast SPGR scan (176 sagittal slices, 1mm^3^ resolution). One anatomical volume was collected per subject. All anatomical volumes were aligned to the anterior commissure - posterior commissure (AC-PC) plane using an affine transformation.

#### Functional

Functional data for pRF mapping and the large-field localizer (see below) were acquired with the 16-channel coil. 36 slices covering occipitotemporal cortex were defined: 2.5mm isotropic voxels, TR 2000ms, TE 29ms, flip angle 77 degree, field-of-view 200×200 mm. Functional data for the small-field localizer (see below) were acquired with the 32-channel coil. 48 slices covering occipitotemporal cortex were defined: 2.4mm isotropic voxels, TR 1000ms, TE 30ms, flip angle 62 degrees, field-of-view 192 x 192mm. An in-plane anatomical image that matched the functional slice prescription was acquired before each set of functional runs. These images were used to align the functional data to the anatomical volume data.

### Stimuli

#### Displays

At the CNI, retinotopic-mapping stimuli were presented with an Eiki LC-WUL100L projector. The Eiki projector has a native resolution of 1920 x 1200 pixels and 10-bit color resolution. Localizer stimuli were either presented with the Eiki projector (large-scale localizer, CNI subjects 13-20) or an LCD display from Resonance Technology (CNI subjects 1-12). The LCD is 47’’ and has a native resolution of 1920 x 1080. Stimuli presented on the LCD display subtended a region of 12 x 12 degrees; stimuli presented with the Eiki projector subtended a region of 32 x 32 degrees.

At the Strauss Center, all stimuli were presented with a NNL LCD display. The display is 32’’ in size and has a native resolution of 1920 x 1080 pixels. Stimuli presented on the display subtended a total region of 12.4 x 12.4 degrees.

#### Functional localizer

At the CNI, stimuli were presented in blocks of eight images from the same category (e.g. words, phase-scrambled objects, etc.) at a rate of 2Hz. Participants performed an odd-ball detection task. Each category was presented six times per run, and three runs were collected per subject, with the localizer experiment lasting roughly 9 minutes. See Le et al. (2017) for more details. At the Strauss Center, stimuli were presented in blocks of six images from the same category (Hebrew text or phase-scrambled objects). Between each text of the phase-scrambled object block was a 10s block of fixation. Each run lasted 3 min 14 seconds, and 2 runs were collected per subject for each language type. Hebrew words were 1 to 6 characters long, with a minimum frequency of 200 per million.

#### Retinotopy

Retinotopy stimuli consisted of bars (4° wide) that slowly and repeatedly traversed the visual field in eight different directions. The pattern within the bar contained checkers, words, small words, false font, or Hebrew words. At the CNI, the width of the bar was 4 degrees. At the Strauss Center, the screen used to display the stimuli is 3.75x smaller; thus the stimuli is also 3.75x smaller (i.e. bar width 1.06 degrees).

Checker stimuli were displayed using code from the Vistadisp software package (https://github.com/vistalab/vistadisp). The checkers within the bars drifted parallel to the orientation of the bar and had a contrast reversal rate of 2Hz. The size of each checker square was approximately 1.3 degree per side. The checker stimuli are described in more detail in previous work (Amano et al., 2009; Dumoulin and Wandell, 2008). Each run was 192 seconds long. Three runs of checker retinotopy were collected for each subject. Four blank periods (12s each) were interleaved to estimate larger pRF sizes to estimate larger pRF sizes (Dumoulin and Wandell, 2008).

All other stimuli types were displayed using code from the analyzePRF software package (http://kendrickkay.net/analyzePRF/). Each sweep of the bar moved continuously across the visual field for 31 seconds for each of the eight orientations. 3 blank periods were included: 16 seconds at the beginning, 16 seconds in the middle, and 20 seconds at the end of each run. The stimuli within the bar (words, small words, false font, Hebrew words) refreshed at a rate of 4Hz.

English words were rendered in black Helvetica font on a white background. The lower-case letter x subtended 1.3 degrees in height. Two-to-six letter words with frequency greater than 200 per million were used as the lexicon; words were randomly chosen from the lexicon to create a page of text. The text within the aperture was refreshed at 4 Hz. Each run lasted 5 min and two runs were collected. The lower-case letter ‘x’ subtended 1.3 degrees in height. For the small word stimuli, the lower-case letter ‘x’ subtended 0.4 degrees in height.

False font and Hebrew words were chosen with the same statistics as the English words (two-to-six letter words, with frequency greater than 200 per million). False font stimuli were rendered in the Futurama font, and Hebrew were rendered in the Arial font. Due to the difference in screen size, Hebrew letters subtended a vertical extent of 0.35 degrees.

### Data analysis

#### Pre-processing

MR data analyses relied on the open-source code in vistalab (https://github.com/vistalab/vistasoft). The basic pre-processing steps included estimation and removal of motion artifacts, and registration of the functional data to the high resolution anatomical images. Motion artifacts within and across runs were corrected using an affine transformation of each volume in a session to the first volume of the first run. In all subjects, head movement was less than 1 voxel (in most cases less than 0.4 voxel). The first 6 time frames of each functional run were discarded. Baseline drifts were removed from the time series by high-pass temporal filtering. The inplane anatomical image was aligned to the average whole brain T1-weighted anatomical image by calculating a rigid body transformation that maximized the mutual information between the inplane anatomy and the resliced volume anatomy. These alignment parameters were then used to align the functional data to the anatomical data.

#### pRF modeling

We model the population receptive field in each voxel using the compressive spatial summation (CSS) model (Kay et al., 2013). The CSS model estimates the pRF of a voxel by selecting parameters for a 2D Gaussian that optimally predicts the response to a translating stimulus (moving bars). The parameters of the pRF include location (x,y) and size (σ) in degrees of visual angle. The value of the 2D Gaussian at its peak is normalized to 1. The CSS model extends earlier linear models by including an estimated compressive nonlinear response exponent to account for subadditive responses. The compression factor increases across the visual hierarchy.

#### Defining the visual field maps and VOTRC

The visual field maps were defined using a probabilistic atlas defined by (Wang et al., 2015). The VOTRC was defined using a combination of functional and anatomical data. The boundaries of the VOT are the inferior temporal sulcus (lateral), the collateral sulcus (medial), hV4 (posterior), and an imaginary horizontal line drawn from the collateral sulcus to the inferior temporal sulcus, starting at the point where the parieto-occipital sulcus would meet the collateral sulcus (anterior). In most subjects, hV4 is found in the posterior transverse collateral sulcus (Witthoft et al., 2014). The selective voxels within this region in each hemisphere are defined as the left or right VOTRC. Selective voxels are more responsive to words than to other visual categories (t-test, p<0.001, uncorrected), but not necessarily contiguous. See (Le et al., 2017) for more details. The VOTRC falls in or near the occipital temporal sulcus in both hemispheres.

## Supplemental Information

**Figure S1.**
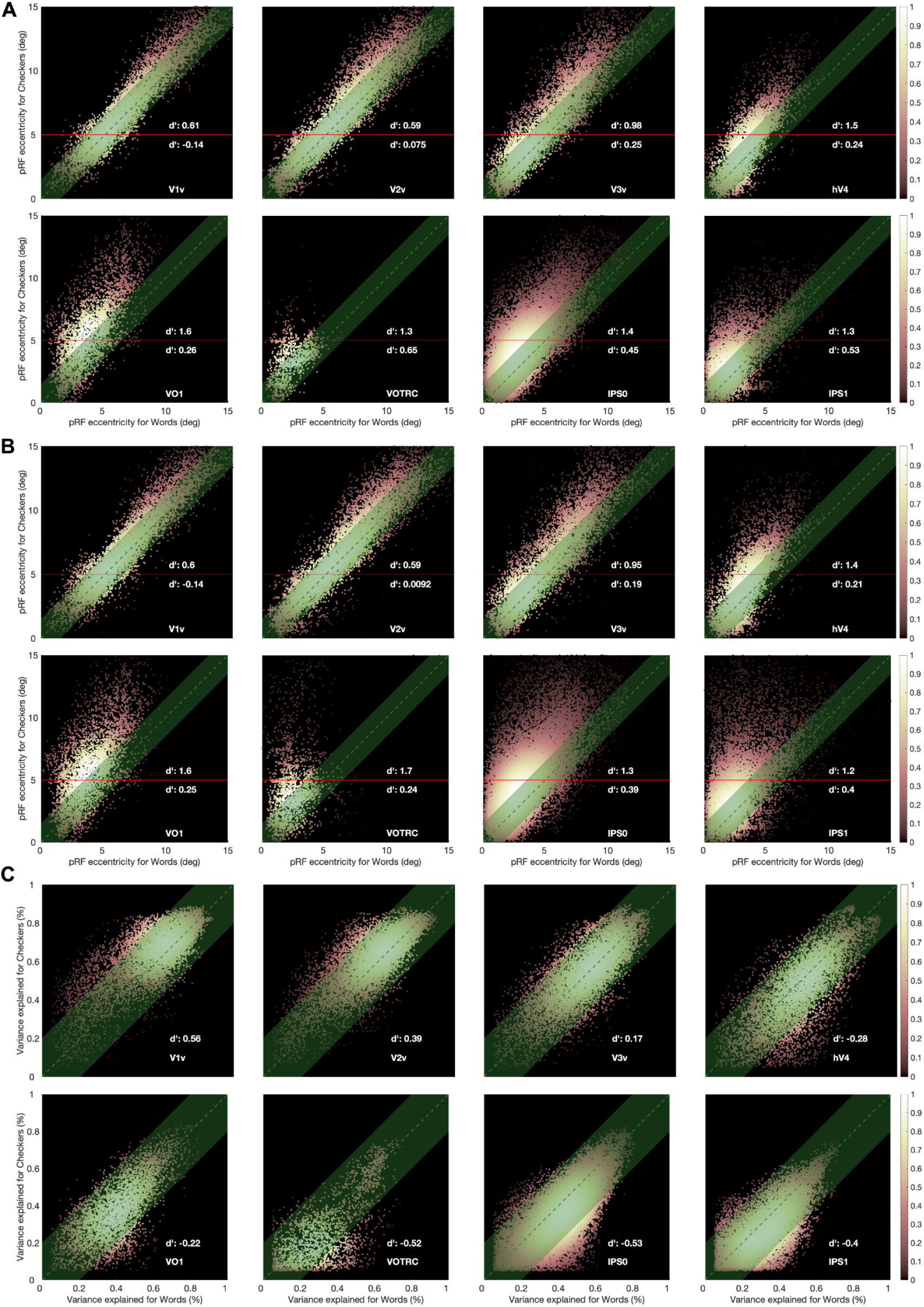
Eccentricity and variance explained for words and checkers (N=20). **A.** Eccentricity for words and checkers restricted to 20% variance explained (VE). **B.** As in A but for 5% VE. **C.** Comparison of VE with words and checkers, restricted to 5% VE or higher. There is a gradual decrease in variance explained from V1 to VOTRC. In V1 VE for checkers is higher than words, and gradually in VOTRC VE for words is higher than checkers.

**Figure S2.**
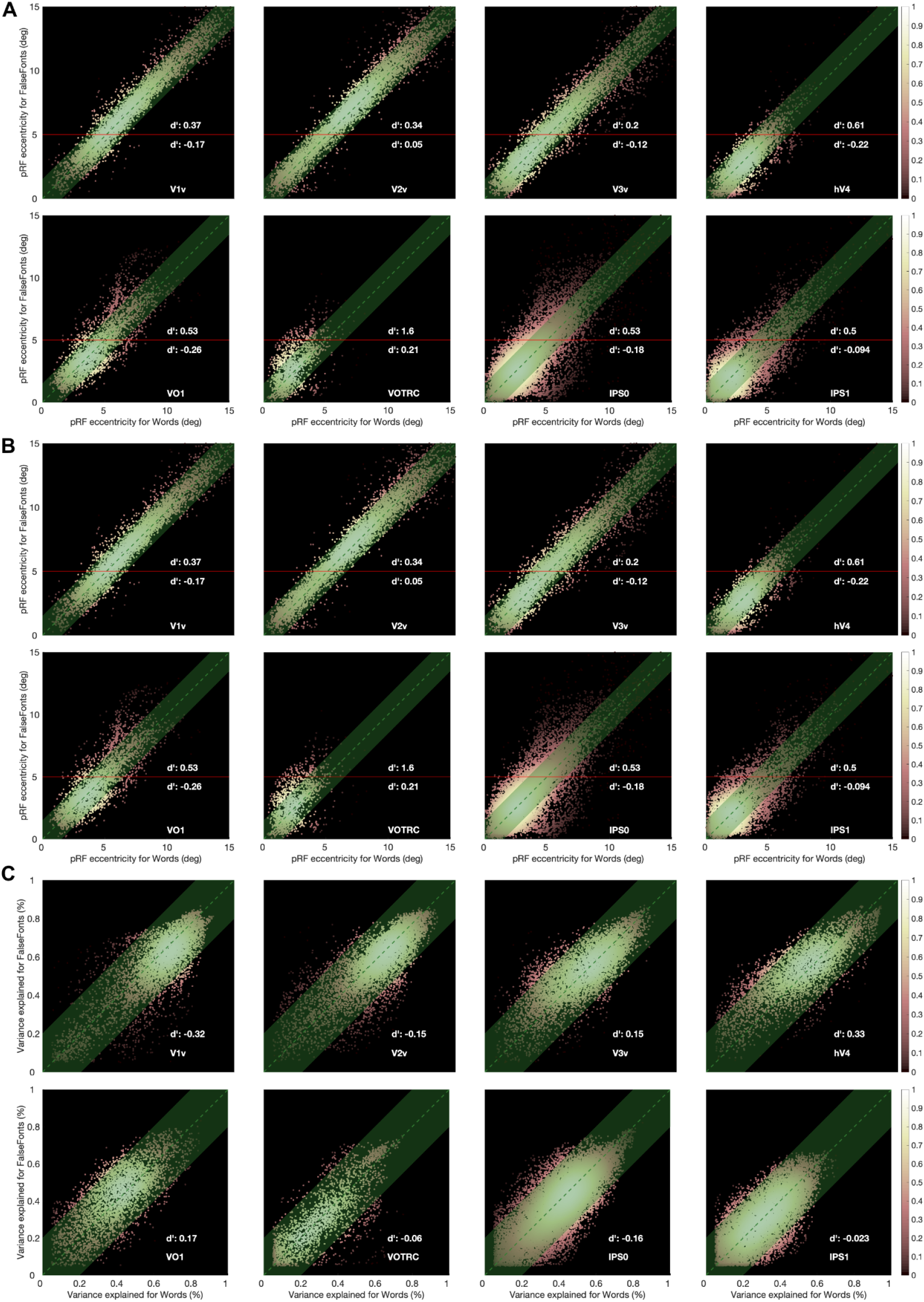
Eccentricity and variance explained (VE) for words and false-fonts (N=20). **A.** Eccentricity for VE at 20%. **B.** Eccentricity for VE at 5%. **C.** VE higher than 5%. VE is similar across all maps. Other details as in Figure S1

**Figure S3.**
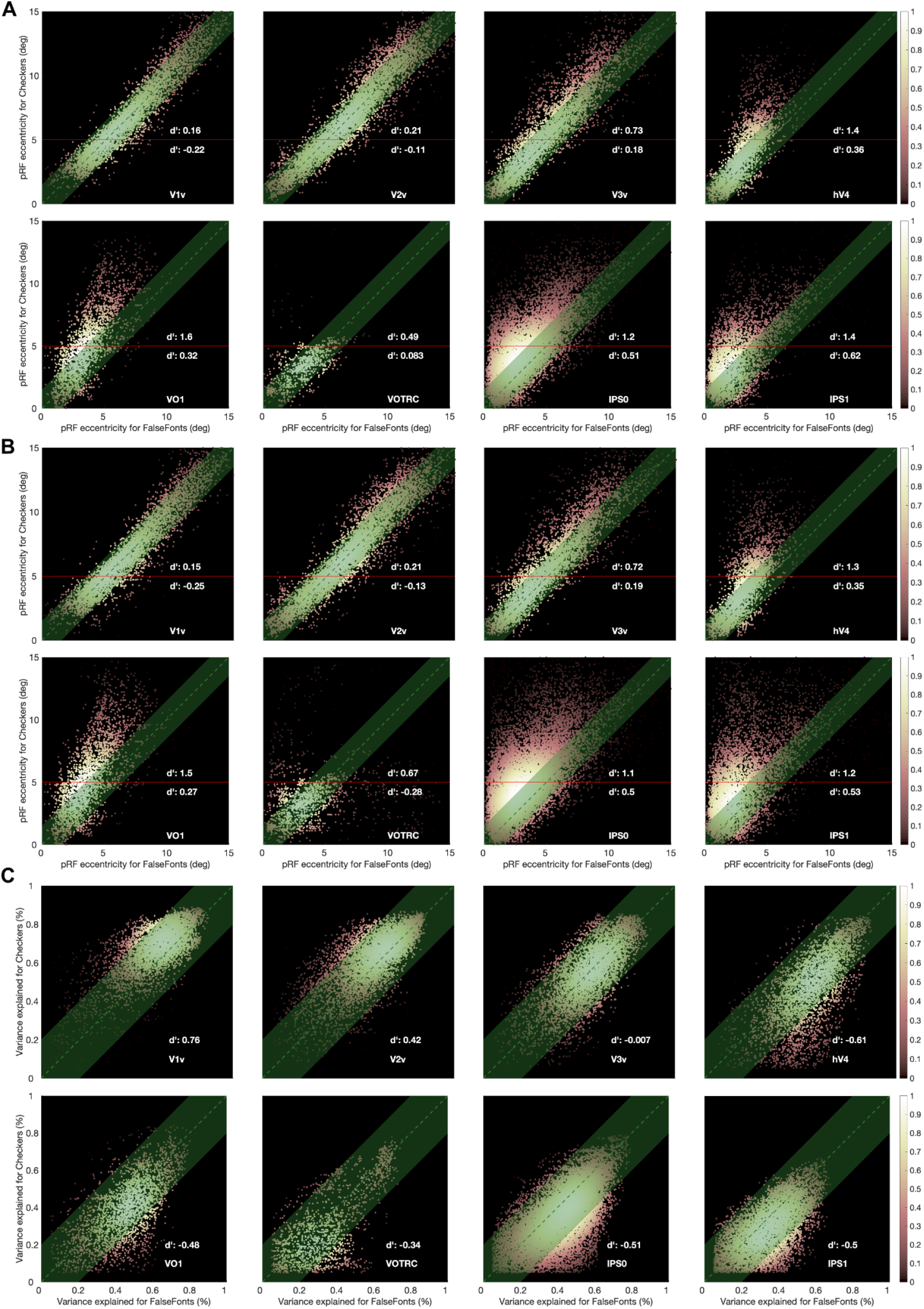
Eccentricity and variance explained (VE) for false-fonts and checkers (N=20). **A.** Eccentricity for VE at 20%. **B.** Eccentricity for VE at 5%. **C.** VE higher than 5%. There is a slight gradual shift in variance explained from V1 to VOTRC. In V1 checkers are higher than false-fonts, and gradually in VOTRC words are higher than checkers.

**Figure S4.**
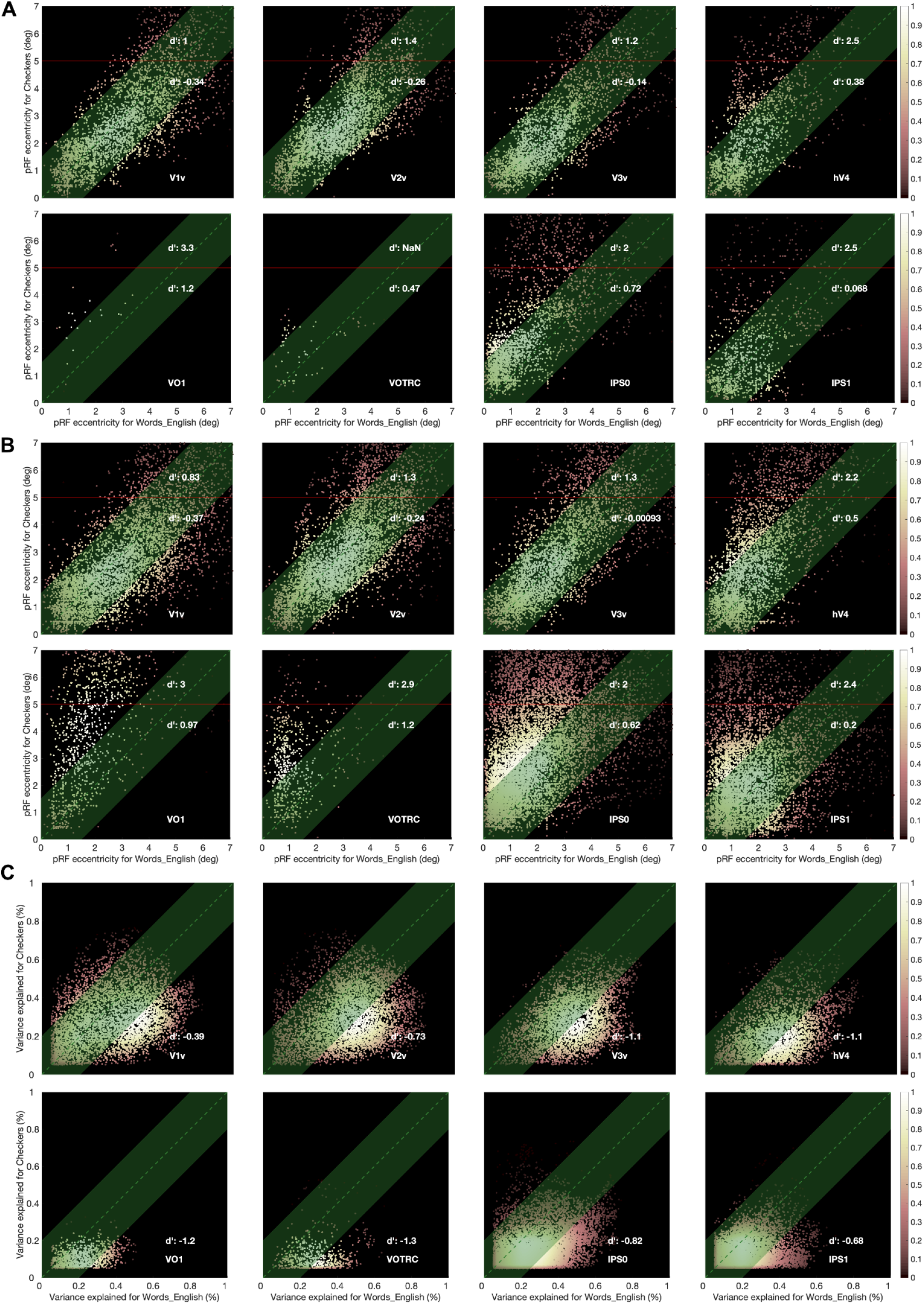
Eccentricity and variance explained (VE) for English Words and Checkers (N=13). **A.** Eccentricity for VE at 20%. **B.** Eccentricity for VE at 5%. **C.** VE higher than 5%. This replicates Fig S1 in another dataset. There is a gradient in VE as well, but it is always higher for Words.

**Figure S5.**
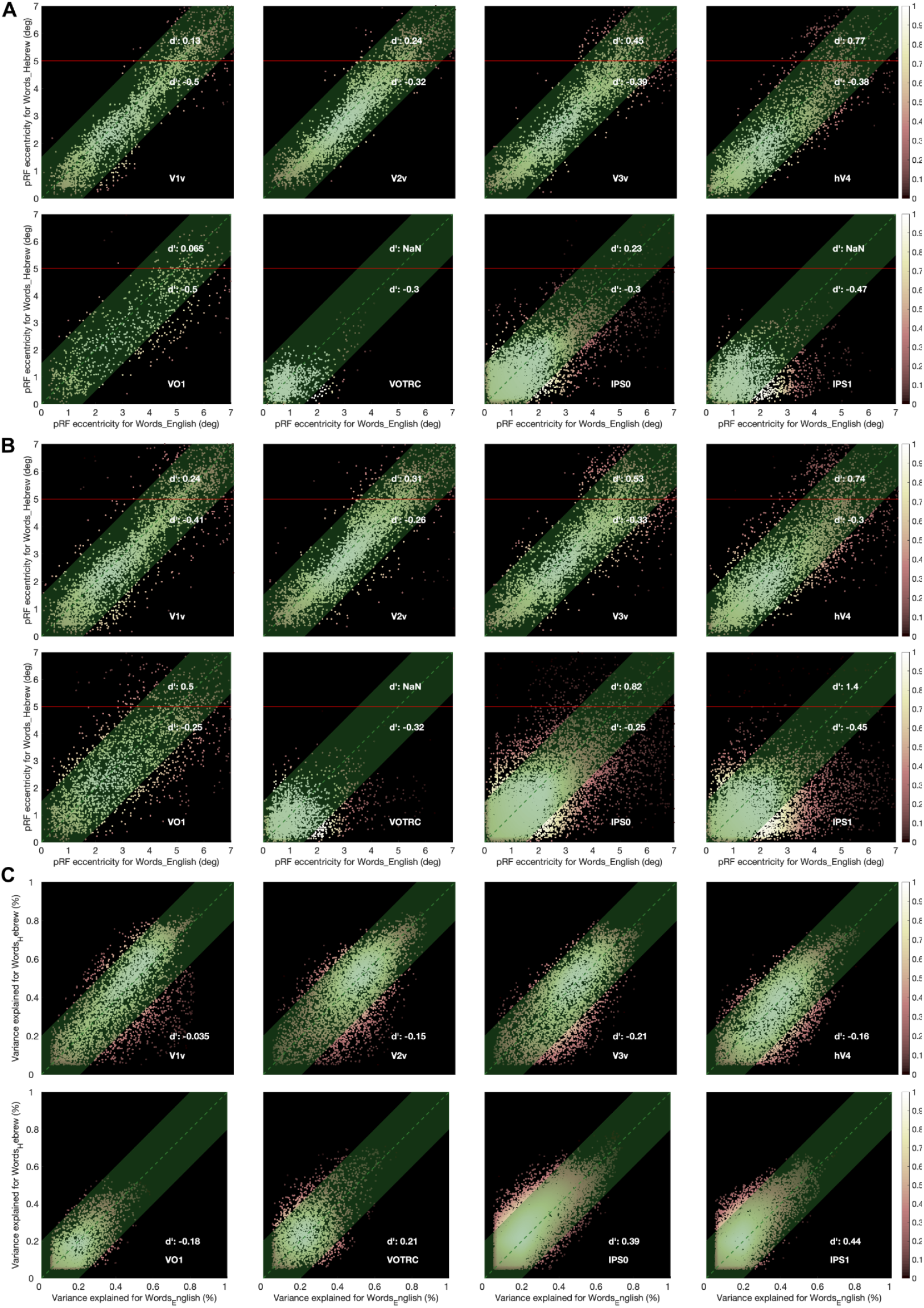
Eccentricity and variance explained (VE) for English Words and Hebrew Words (N=13). **A.** Eccentricity for VE at 20%. **B.** Eccentricity for VE at 5%. **C.** VE higher than 5%. VE is similar across all maps.

**Figure S6.**
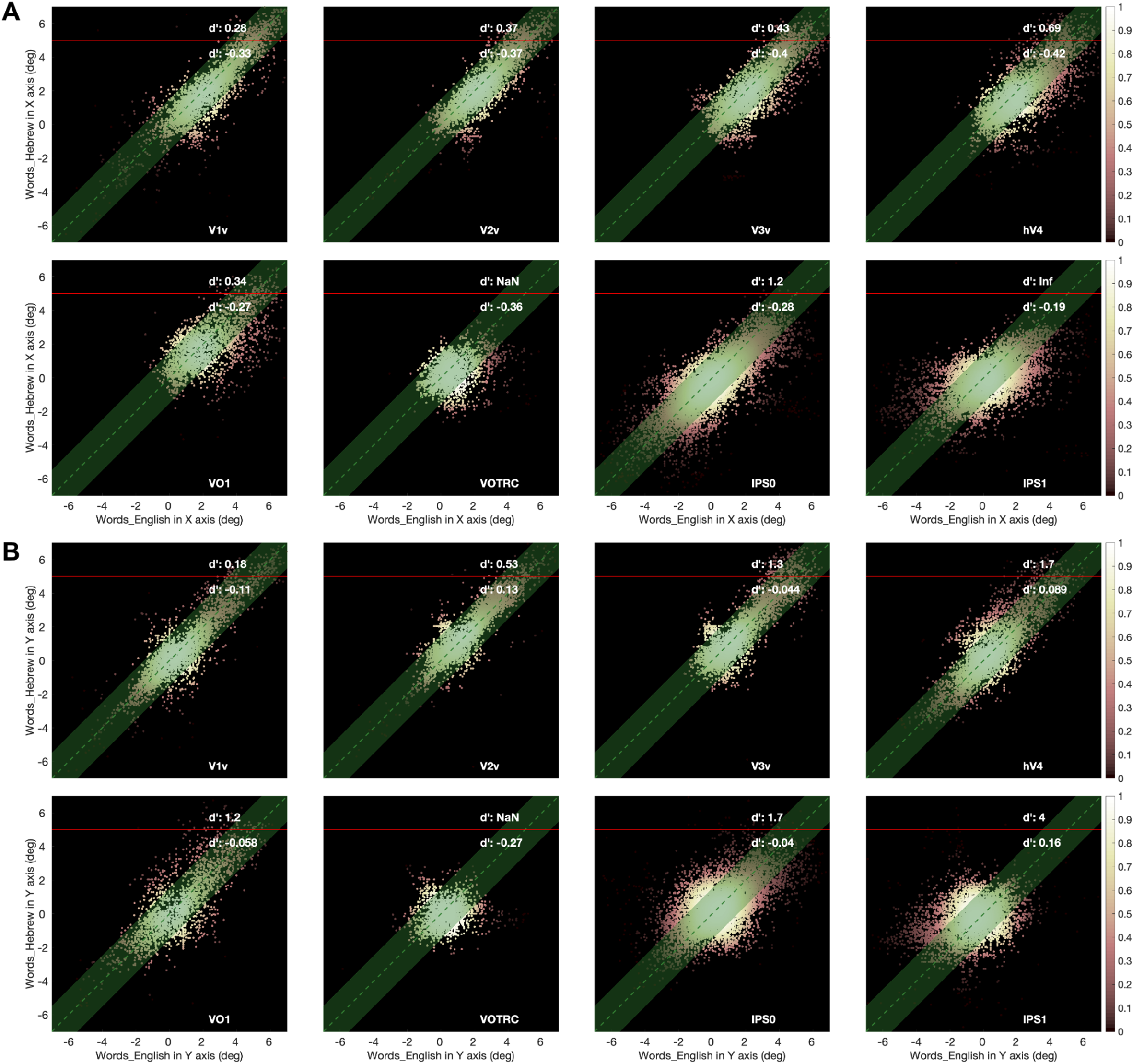
X and Y coordinates of pRF center estimates for English Words and Hebrew Words (N=13). **A.** X coordinate at VE 5%. **B.** Y coordinate at VE 5%. T-test values per each region (English Words - Hebrew Words): **V1v_left** (N=3010): (X) P:3.04183e-104, CI: [−0.288887 −0.242685]; (Y) P:0.000553134, CI: [−0.0614414 −0.0169712] **V2v_left** (N=4095): (X) P:3.84073e-93, CI: [−0.194706 −0.161467]; (Y) P:6.39918e-18, CI: [0.0538268 0.085307] **V3v_left** (N=3634): (X) P:6.92128e-152, CI: [−0.31778 −0.275553]; (Y) P:0.00698734, CI: [0.00736248 0.0464623] **hV4_left** (N=3623): (X) P:4.72068e-104, CI: [−0.294123 −0.246748]; (Y) P:0.651941, CI: [−0.0277358 0.01736] **VO1_left** (N=874): (X) P:6.29476e-38, CI: [−0.526074 −0.392648]; (Y) P:2.20269e-09, CI: [−0.25777 −0.131422] **lVOTRC** (N=1334): (X) P:8e-36, CI: [−0.274353 −0.201787]; (Y) P:1.37949e-57, CI: [−0.33622 −0.265901] **IPS0** (N=6996): (X) P:3.30683e-157, CI: [−0.282283 −0.244602]; (Y) P:1.65323e-58, CI: [−0.191547 −0.150372] **IPS1** (N=3437): (X) P:1.40678e-46, CI: [−0.27722 −0.211374]; (Y) P:3.926e-07, CI: [0.0488939 0.110311]

**Figure S7.**
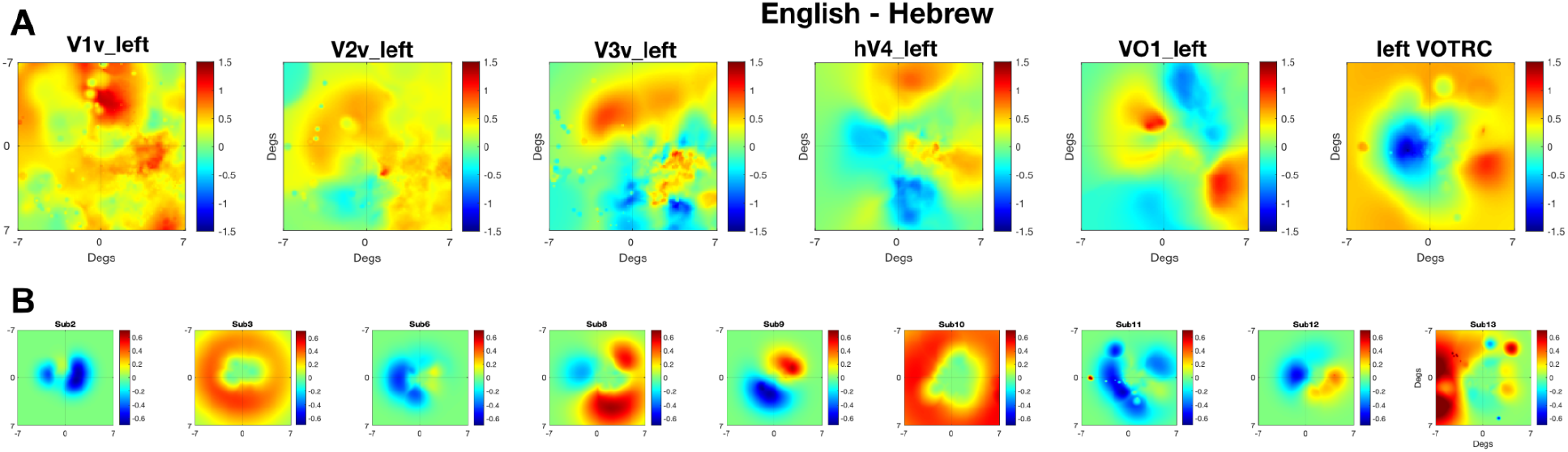
FOV differences between English Words and Hebrew Words. **A.** English - Hebrew FOV differences (Cohen’s d’) in six different regions. VE at 20%. N = 9. **B.** English - Hebrew FOV individual subject differences. VE at 20%.

